# Prey identity affects fitness of a generalist consumer in a brown food web

**DOI:** 10.1101/2021.09.02.458769

**Authors:** Lily Khadempour, Leslie Rivas Quijano, Casey terHorst

**Affiliations:** Department of Biology, California State University, Northridge, Northridge, CA, 91330, USA; Department of Earth and Environmental Sciences, Rutgers University, Newark, Newark, NJ, 07102, USA

**Keywords:** *Sarrecenia purpurea*, phytotelmic, predator-prey dynamics, bottom-up effects

## Abstract

The use of ever-advancing sequencing technologies has revealed incredible biodiversity at the microbial scale, and yet we know little about the ecological interactions in these communities. For example, in the phytotelmic community found in the purple pitcher plant, *Sarrecenia purpurea*, ecologists typically consider the bacteria as a functionally homogenous group. In this food web, bacteria decompose detritus and are consumed by protozoa that are considered generalist consumers. Here we tested whether a generalist consumer benefits from all bacteria equally. We isolated and identified 22 strains of bacteria, belonging to six genera, from *S. purpurea* plants. We grew the protozoa, *Tetrahymena* sp. with single isolates and strain mixtures of bacteria and measured *Tetrahymena* fitness. We found that different bacterial strains had different effects on protozoan fitness, both in isolation and in mixture. Our results demonstrate that not accounting for composition of prey communities may affect the predicted outcome of predator-prey interactions.

## Introduction

Predator-prey or consumer-resource dynamics are among the best studied ecological interactions. Theory suggests that coexistence of predator and prey depends on several parameters that incorporate traits of both predator and prey, such as attack rate, handling time, and conversion efficiency (Holling 1959). Considerable theory has explored how variability of predator and prey traits affects population dynamics, demography, and species coexistence (e.g. Abrams and Rowe 1996; Kendall et al. 1999; Peckarsky et al. 2008; Fleischer et al. 2018). Many empirical examples demonstrate how different prey traits affect handling time (e.g. Werner 1974; Faria et al. 2004) and attack rates of predators (Boates and Goss-Custard 1992; Elliott 2003) ultimately having consequential effects on predator fitness. It would come as no surprise to many ecologists that the identity of an animal’s prey affects performance and fitness, yet in the microbial world, we have far less understanding of the specificity of trophic interactions.

Most examples of trophic interactions emerge from green food webs, where photosynthetic plants or algae form the base of the food web. However, brown food webs, in which detritivores, such as bacteria and fungi, form the base of the food web, are also common in nature. In green food webs, ecologists are keenly aware of how traits of the plant community affect the transfer of energy through a food web (Stap et al. 2007; Mooney et al. 2010). However, in brown food webs, the species composition of the base of the food web has historically been treated as a black box where “bacteria” or “fungi” are treated as a single taxonomic unit (eg. Miller and terHorst 2012). Despite rapid advances in identifying bacterial taxa in natural communities over the past decade, we still have far less knowledge of the ecological role of specific taxa and how they interact with other species. The effect of different bacterial species on consumer growth rates has remained largely untested (but see Mohapatra and Fukami 2005; Darby and Herman 2014), even though variation in bacterial species traits are likely to alter consumer attack rates, handling times, and conversion efficiencies.

The phytotelmic (organisms that inhabit small pools of water within or upon plants) community found in the leaves of the purple pitcher plant (*Sarrecenia purpurea*) has been used as a model system for studying broad questions about ecology and evolution (Miller et al. 1994; Cochran-Stafira and Ende 1998; Ellison and Gotelli 2002; Kneitel and Miller 2002; Miller et al. 2014). The pitcher-shaped leaves attract insects and serve as pitfall traps in which insects drown, decompose, and provide nutrients to the plant. The insects serve as the source of energy and nutrients at the bottom of a brown food web. Bacteria decompose the dead insects and are consumed by a suite of protozoa and rotifer species, which are consumed by mosquito larvae. Numerous top-down studies have demonstrated that the evolutionary and ecological dynamics of protozoan consumers affect bacterial community composition (Cochran-Stafira and Ende 1998; Peterson et al. 2008; Paisie et al. 2014; Holdridge et al. 2016; Canter et al. 2018). These top-down effects should not be surprising since protozoans have demonstrated selective feeding behavior in other systems (Strom and Loukos 1998; Gaines et al. 2019). Selective feeding clearly has an effect on prey community dynamics, and it also suggests that different prey could have different effects on consumer fitness. However, there is considerably less work on the bottom-up effects of different bacterial species on protozoan ecological dynamics. Although some studies have examined the effects of total bacterial abundance on higher trophic levels (e.g., Kneitel and Miller 2002; Hoekman 2007), most studies indirectly manipulate the bacterial community as a whole by altering resource availability, rather than particular taxa of bacteria with different traits. An underlying assumption with these studies is that the protozoa are generalist consumers of bacteria and that, regardless of their identity, a higher abundance of bacteria promotes protozoan growth.

Here we examine this assumption and ask whether different strains of bacteria differentially affect consumer fitness. We collected fluid from *S. purpurea* pitcher plants found in the field, from which we isolated single strains of bacteria. We quantified the effects of single strains and multi-strain communities of bacteria on the fitness of a common ciliate (*Tetrahymena* sp.), which is found in *S. purpurea* phytotelmic communities, and is commonly used in lab microcosm experiments in this system.

## Methods

### Isolation of bacteria

We collected water contained within pitcher plant leaves haphazardly from leaves of various ages in one field in the Apalachicola National Forest in northern Florida (USA). Large insect parts were filtered from the fluid and 2% v/v DMSO was added before freezing at −20°C in 50 mL conical tubes. Samples were shipped to California State University, Northridge, where we thawed the tubes and diluted the liquid with sterilized water at 10×, 100×, and 1000× dilutions. We spread 50 *μ*L of liquid from each dilution onto LB solid media plates (Cold Spring Harbour Protocols 2016) using a plate spreader. We monitored these plates daily to check for bacterial colony growth and then identified different morphotypes of bacteria, which were picked and streaked in order to isolate individual strains. Once we were confident the bacteria were in isolation, both through visual confirmation and sequencing (see below), we maintained them in 5 mL of liquid LB media (Cold Spring Harbour Protocols 2016) and transferred them to fresh tubes every two weeks.

### Identification of bacteria

We extracted DNA from bacterial strains using the Qiagen Blood and Tissue Kit, with the specifications for bacterial cultures. We used 16S rRNA gene primers 27F and 1392R in a PCR with the following conditions: 94°C for 5 min, 30 cycles of 94°C for 20 s, 55°C for 20 s, and 72°C for 70 s, with a final elongation of 72°C for 10 min. We sequenced the amplicons with Sanger sequencing through Laragen (Culver City, CA) with both forward and reverse primers, then trimmed, identified, and analyzed sequences using 4Peaks and CLC Sequence Viewer 7. We used NCBI BLAST to identify the strains to the genus level using their 16S sequences. Ultimately, we isolated 22 individual unique strains of bacteria (Table S1).

### Isolation of protozoa

For several years, we have maintained eight strains of the ciliated protozoa *Tetrahymena* sp. in lab cultures. These strains were originally collected from different pitcher plants in the Apalachicola National Forest and have been maintained independently in the lab. For this experiment, we isolated each of these strains from their associated bacterial community. We created YPD sterile media (YPD media 2010) supplemented with four filter-sterilized antibiotics (final concentration in parentheses): kanamycin (50 mg/mL), tetracycline (10 mg/mL), penicillin (1200 mg/mL), and streptomycin (120 mg/mL). We added 100 *μ*L of each *Tetrahymena* strain to separate replicate test tubes with media and allowed them to grow for three days at room temperature. We then looked for bacteria in a small volume of the protist culture at 1000× magnification. We only used replicate tubes in which little or no bacteria were visible. This technique was unlikely to have removed all bacteria, but the concentration of bacteria in these cultures was many orders of magnitude lower than the bacteria that we added in the experimental tubes below. We then added each protist strain to the experimental tubes, as described below.

### Effect of individual bacterial strains

We established microcosms by adding 10 mg of crushed Tetracolor Fish Flakes to 10 mL of water in a test tube before autoclaving for 45 min at 120 °C. We added 30 *μ*L of liquid culture of an individual bacterial strain to each tube and allowed these bacteria to establish and grow for one day at ambient room temperature. We did not control for bacterial abundance as we considered bacterial growth rate on the media as one of several bacterial traits that could potentially differ among strains. We then added an individual *Tetrahymena* strain to different tubes, so that each *Tetrahymena* strain was grown with each bacterial strain; each *Tetrahymena* strain served as a replicate (n=8) in testing the effects of the bacterial strains. The initial density of protozoa in each tube was 100 individuals per mL. We allowed the bacteria and protozoa to grow together for four days at ambient room temperature, and then counted the density of protozoa using Palmer cells (Wildlife Supply Company, Yulee, Florida, USA).

### Effect of multiple bacterial strains

To determine whether different bacterial strains would have different effects when combined with other bacterial strains in a community context, we also created synthetic communities of bacteria. We created five different combinations of six bacterial strains. From the first experiment, we chose (1) the six best strains, in terms of their effect on protozoan fitness, (2) the six worst strains, (3) the three best and three worst strains, (4) the two best, two worst, and two intermediate strains, and (5) a combination of the six worst strains that had different morphotypes (Table S2). We grew each of these five bacterial combinations with each of the eight unique *Tetrahymena* strains. We mixed equal volumes of the six strains in each combination and then aliquoted 30 *μ*L of this mixture into each replicate experimental tube. We allowed these bacterial cultures to grow for one day before adding protozoa and quantifying their growth in the same way as in the previous experiment.

For both experiments, we used ANOVA to examine the effects of bacterial strain or mixture on the final abundance of protozoa. *Tetrahymena* strains were considered as independent replicates. We followed significant treatment effects with Tukey’s HSD to examine pairwise differences among treatment levels, using the ‘agricolae’ package (Mendiburu and Yaseen 2020) in R 3.6.3 (R Core Team 2013).

## Results and Discussion

In total, we isolated 22 bacterial strains with different morphotypes, belonging to six genera: *Serratia, Chromobacterium, Chryseobacterium, Burkholderia, Acinetobacter*, and *Bacillus*. Individual strains of bacteria had different effects on protozoan fitness (F_(21,153)_ = 14.2, P < 0.001); Figure 1). Some bacterial taxa resulted in highly abundant protozoan populations, although other taxa either could not support protozoan growth or actively inhibited it, and many other taxa had varying degrees of intermediate effect.

**Figure 1.**
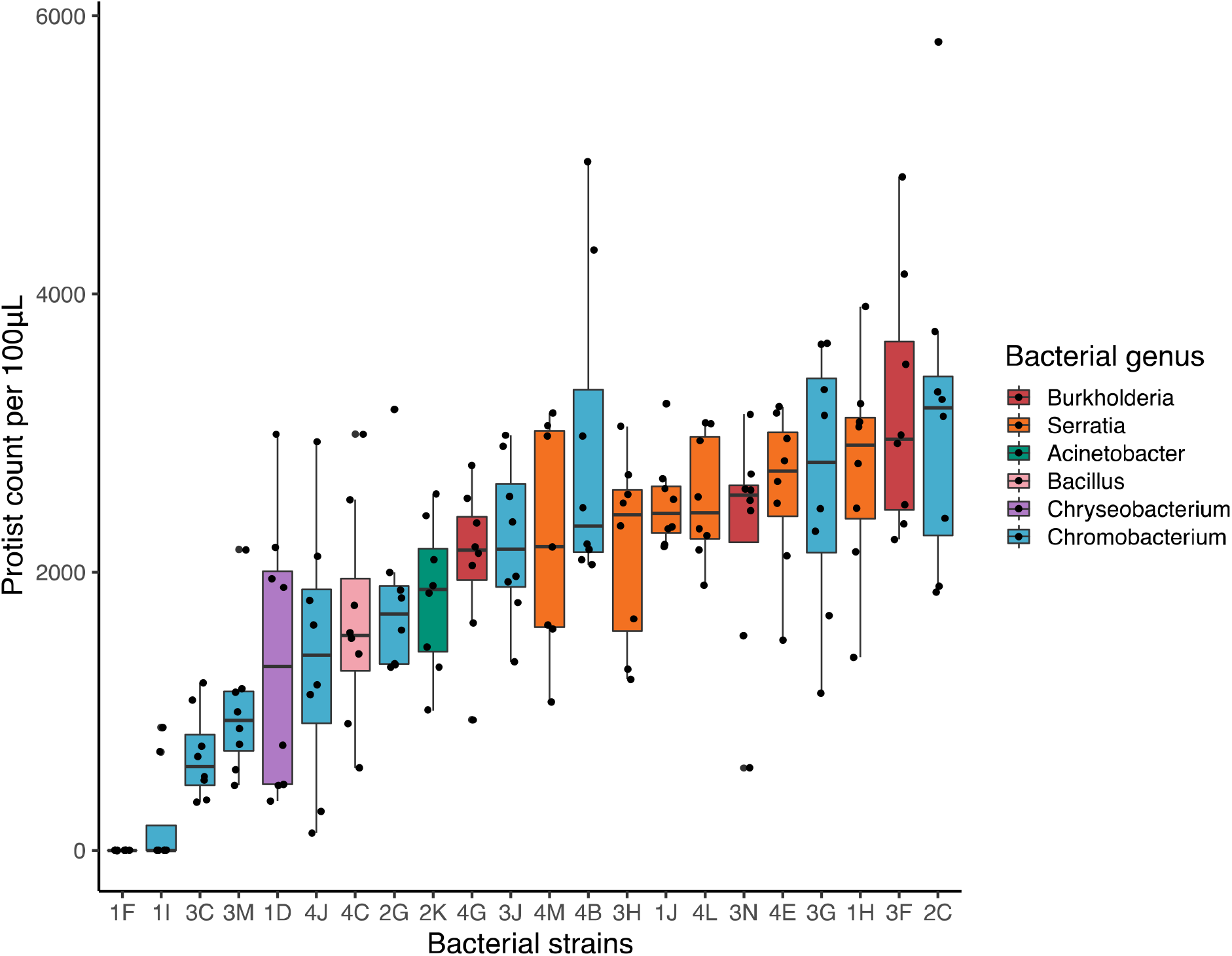
Protist abundance depends on the identity of bacterial strain (F_(21,153)_ = 14.23, P < 0.001). Box plot definitions: center line – median, upper and lower box limits – upper and lower quartiles, whiskers – 1.5 × inter-quartile range, outliers – any points outside the 1.5× inter-quartile range.

If *Tetrahymena* is indeed a generalist consumer with regard to what it consumes, then it is decidedly not a generalist with regard to the benefits it receives from consuming prey. Instead, bacterial taxa reside on a broad spectrum in regard to their effects on protozoan fitness. Although many studies in this system assume that protozoan fitness is primarily determined by the abundance of bacterial prey, our results suggest that fitness may be largely determined by the composition of the bacterial community.

Two *Chromobacterium* strains (1F and 1I) had especially negative effects on protozoan fitness, with nearly no protozoa observed in these cultures. This is consistent with previous studies that have shown that *Chromobacterium* spp. inhibit protozoan growth through the production of the pigment violacein (Singh 1945; Pickup et al. 2007). For this reason, *Chromobacterium* has been targeted as a potential source of compounds for treating fungal and viral infections, as well as cancer cell growth (Cheng et al. 2007; Sasidharan et al. 2015). However, other strains of *Chromobacterium* (4B, 3G, and 2C) produced some of the highest protozoan abundances. This emphasizes that the ecological function of bacteria cannot be generalized, even among closely-related taxa; single strains of bacteria may not well represent the effects of other species in the same genus. Other genera, such as *Burkholderia* and *Serratia*, had consistently positive effects on protozoan fitness, although more isolation and testing of other taxa within these genera are necessary to determine if this is generally true. The effects on bacteria traits on protozoa growth are not limited to only toxicity, as there were a range of intermediate effects of different bacteria taxa. Other factors such as, but not limited to, growth rate, motility, and nutritive quality might also affect protozoan fitness, and a more detailed analysis of bacterial traits and their effects on protozoans would provide for interesting future work (Goyal et al. 2021).

Our pairwise interaction experiment demonstrated that bacterial effects on protozoan fitness depend on bacterial strain identity. However, in natural communities, protozoa are unlikely to interact with one bacterial species in isolation. In our second experiment, we found that different community compositions had different effects on protozoan abundance (F_(4,34)_ = 29.5, P < 0.001; Figure 2). The negative effects of the “bad” bacterial strains (those that had strong detrimental effects on protozoan fitness) persisted in a community context, where the effects of bad strains outweighed the positive effects of relatively “good” strains (those that have relatively beneficial effects on protozoan fitness). Any combination that included the two bad bacteria from the first experiment (1F and 1I) resulted in low protozoan fitness (Figure 2). This pattern is consistent with bad strains having produced toxins with effects that were not dampened by the presence of other species, rather than the bad strains not supplying sufficient nutrition for the protozoa. It is important to note that our synthetic communities only consisted of six bacterial strains, which is considerably less diverse than a natural community found in a pitcher plant leaf (Koopman et al. 2010; Koopman and Carstens 2011; Gray et al. 2012), reported to contain approximately 400 bacterial species (Paisie et al. 2014). It is unclear whether the negative effects of the bad bacteria would be dampened in a more diverse bacterial community in a pitcher plant leaf, which has many more microhabitats than a test tube, and where the bad bacterial species would face greater and more diffuse competition for resources with other bacteria.

**Figure 2.**
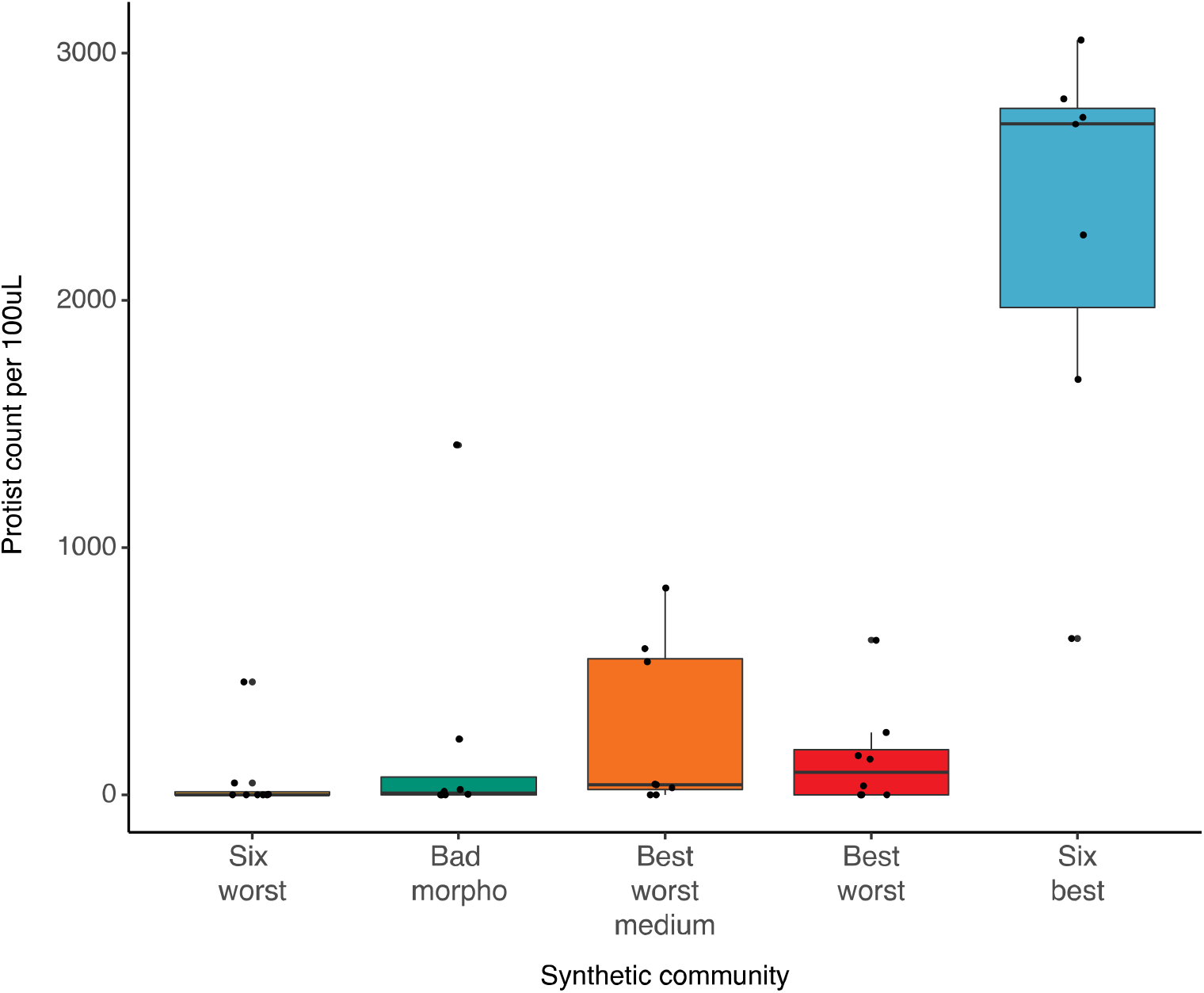
Protist abundance varies depending on a mixture of strains (F_(4,34)_ = 29.46, P < 0.001). Any strain combination that contained harmful strains of *Chromobacterium* resulted in poor protozoan growth. Box plot definitions: center line – median, upper and lower box limits – upper and lower quartiles, whiskers – 1.5 × inter-quartile range, outliers – any points outside the 1.5× inter-quartile range.

Simplistically, bacteria are thought to be beneficial to the pitcher plant because they break down insects and release nutrients, while protozoa are thought to be parasitic because they consume bacteria (Mouquet et al. 2008). Our results suggest that these categorizations may be more complex since protozoa do not necessarily reduce the abundance of all bacterial taxa evenly. This result also supports previous work in this system and others that demonstrates that protozoa have different prey selection patterns (Strom and Loukos 1998; Gaines et al. 2019). Furthermore, presumably some bacterial taxa break down prey faster than others, although this has not been tested in this system, to our knowledge, nor do we know whether there is a trade-off between this function and susceptibility to predation. We also do not yet know whether the same taxa-specific effects of bacteria affect other protozoan consumers in the same way. Future work could expand on these results by doing similar manipulations inside pitcher plant leaves and examining insect degradation rates or nutrient uptake by the plant. Such information would help us to understand how dynamics among species within the phytotelmic community ultimately affect the plant in which they live.

When *Tetrahymena* protozoa feed, they appear to ingest any bacteria in their immediate surroundings, although they often cluster around structures in the water (e.g., insect parts or pieces of detritus; personal observation). The extent to which bacteria growing on structures in the water differ from those in the water column, or whether good and bad bacteria differ between these environments is unknown and would be an interesting avenue for future work in this system. It has been established that protozoa have top-down effects on bacterial community composition (Cochran-Stafira and Ende 1998; Peterson et al. 2008; Paisie et al. 2014; Holdridge et al. 2016; Canter et al. 2018), and this, combined with the knowledge that different strains affect protozoan fitness, would suggest that the protozoa may engage in a more active form of prey selection than previously thought. There are, however, alternative explanations for the top-down effects and prey selection patterns of the protozoa on bacterial community composition. For example, some bacterial taxa may have faster growth rates, and so can recover more quickly from grazing, or some bacteria may congregate in microniches where protozoa are more abundant, making them more likely to be eaten.

Overall, our results demonstrate that bacterial prey identity affects consumer fitness and that it is vital that future work in these pitcher plant communities account for bottom-up effects of different bacterial taxa on protozoan growth and fitness, both by taking this into account in models, but also by considering this in empirical studies. For example, Miller et al. (2012) found only a weak relationship between bacterial abundance and protozoan abundance across a successional sequence in pitcher plant leaves. However, our work suggests that examining relationships between particular types of bacteria and their effects on the abundance of different protozoa species, and vice versa, could reveal previously unrecognized ecological patterns. Future work could consider how to manipulate or quantify the bacterial community in such a way to tease apart these dynamics. These results open up new avenues of research in pitcher plant microcosms that allow for the study of bottom-up effects by maintaining lab protozoa aseptically, or with known bacterial taxa, allowing for manipulation of the bacterial community during lab experiments. Alternatively, researchers who either create their own synthetic communities or identify bacterial community members through shotgun sequencing can be aware of which bacterial taxa are present and can account for those effects appropriately, rather than treating the bacterial community as a single unit.

Like other brown food webs, pitcher plant microbial communities are, in practice, often treated as food chains, where the bacteria at the base of the food web are assumed to be functionally redundant. This study demonstrates that this assumption is not valid, and that when bottom-up effects are being examined in this system, it is important to consider the identity of the bacterial community members and their effects on the predators that feed on them. We believe that the results of this study can be further extrapolated to other brown food webs, suggesting that we must move away from treating the foundation of these webs as a black box.

## Acknowledgements

The authors would like to thank T. Miller and C. Cuellar-Gempeler for their help in collecting bacteria from the field and members of the CSUN EcoEvo Lab for their feedback on an earlier version of the manuscript. This project was funded by grants from the National Science Foundation to CPT (OCE-1559105 and DEB-1754449).

## Author contribution statement

Conceived of the idea: LK and CPT. Conducted the experiment: LK and LRQ. Wrote the manuscript: LK, LRQ, and CPT.

## Notes

### Competing Interest Statement

The authors have declared no competing interest.

